# A Bayesian inference tool for identifying artifactual calls from differential transcript abundance analyses

**DOI:** 10.1101/2020.02.27.967240

**Authors:** Stefano Mangiola, Evan A Thomas, Martin Modrák, Anthony T Papenfuss

**Author notes:** Correspondence: Stefano Mangiola, Anthony T Papenfuss.

## Abstract

Relative transcript abundance has proven to be a valuable tool for inferring the phenotype of biological systems from genetic material. Several methods for the analysis of differential transcript abundance have been developed, and some of the most popular are based on negative binomial models. Although most genes are fitted reasonably well by the negative binomial distribution, the presence of outlier observations that do not fit such models can lead to artifactual identification of significant changes in transcription. Identifying those transcripts for the correct interpretation of results is extremely important. A robust and automated tool for detecting sample/transcript pairs that do not fit a negative binomial regression model is currently lacking. Here we propose ppcseq, a robust statistical framework that models hierarchically sample- and gene-wise features such as sequencing depth bias, the association between mean transcript abundance and its over-dispersion, and provides a theoretical transcript abundance distribution, on which the observed transcript abundance can be tested for outliers. We show using a publicly available data set where nearly 10% of differentially abundant transcripts had fold change inflated by the presence of outliers. This method has broad utility in filtering artifactual results of differential transcript abundance analyses based on a negative binomial framework.

## Introduction

The analysis of the relative abundance of transcript copies is valuable for inferring the phenotype of biological systems. The sequencing of RNA molecules involves sampling from the population of transcripts present in solution at the time of RNA extraction; this number reflects the relative proportion/concentration of each transcript. Some of the most popular software for differential transcript abundance analysis (e.g. edgeR (Robinson *et al.*, 2010), DeSeq2 (Love *et al.*, 2014)) model the transcript abundance using a negative binomial numerical process. The negative binomial distribution has independent parameters for mean and over-dispersion, and can be thought as an extension of the Poisson distribution, where the mean parameter is generated from a gamma distribution. The negative binomial distribution can be interpreted as a model of two types of variability: (i) the biological variability in mRNA synthesis/degradation rates between replicates (the gamma distribution) and (ii) the intrinsic variability in mRNA counts given constant synthesis/degradation rate and the inherently imperfect efficiency of mRNA extraction and sequencing (the Poisson distribution). Previous studies demonstrated that transcript abundance across samples is characterised by a quadratic relationship between its mean and its variance (Van den Berge *et al.*, 2019), on which several differential transcript abundance algorithms are based, such as DeSeq2 (Love *et al.*, 2014) and edgeR (Robinson *et al.*, 2010). Although most genes are reasonably well-fitted by the negative binomial distribution, the underlying gamma distribution has relatively thin tails and thus is not robust against the presence of unmodelled large-scale biological variability. Larger than expected variability results in some samples (outliers) having disproportionate influence on the final inference, increasing both false positives and false negatives. The attention that several popular methods (Love *et al.*, 2014; Liu *et al.*, 2015) give to outlier detection provides evidence for the importance of the matter.

Although the analysis of errors between the inferred theoretical distribution and the data (i.e., residuals) is possible, this relies on a sufficiently large sample size and would require care in order to consider the information about overall uncertainty of the inferred model. A robust and automated tool for detecting sample/transcript pairs that are outside a negative binomial regression model is currently missing. Bayesian inference provides a robust methodology to simulate the theoretical data distribution according to the inferred model, which includes the integrated uncertainty of the hierarchical parameters (i.e. a posterior predictive check), and therefore is suitable for low-data regimes. The observed data can be mapped against the theoretical data distribution and posterior quantiles of the observed data points can be computed. If those quantiles are close to extremes (0 or 1), it indicates there is a possible mismatch between the model and the data. Furthermore, with the Bayesian inference framework it is possible re-fit the model omitting the suspected outlier data-points, resulting in a more conservative test of the negative binomial assumption. Here we describe ppcseq, an algorithm that is able to (i) model RNA sequencing transcript abundance using hierarchical negative binomial regression; (ii) produce theoretical data distribution with and without possible outliers; and (iii) provide evidence in terms of probability that specific data points do not follow the negative binomial assumption. On publicly available data we identified 9.5% of transcripts with fold change inflated by the presence of outliers, including the second most highly ranked transcript.

## Methods

### Iterative outlier detection

In order to identify the transcripts that partially violate the negative binomial assumption, three types of uncertainty are modelled jointly from the data (Fig. 1): (i) the mean abundance and overdispersion of transcripts, and their associations; (ii) the effect of sequencing depth; and (iii) the association between transcript abundance and the factors of interest. The inference workflow consists of two iterative steps (Fig. 2); potential outliers are identified in a first *discovery* step, and a probability estimation is given on a model excluding those data points in a *test* step. The motivation is twofold. First, after some outliers have been identified, the model needs to be refit as those outliers might have skewed the initial estimates noticeably. In theory this process would need to be iterated until convergence, but in practice we found using two iterations was satisfactory. Second, the stringency of the check for outliers can be set separately for each step. That is, we can identify potential outliers with a loose criteria, refit the model and then check whether those points remain outliers against the refitted, more robust model but with more stringent criteria, letting us improve both sensitivity and specificity of the method.

**Figure 1.**
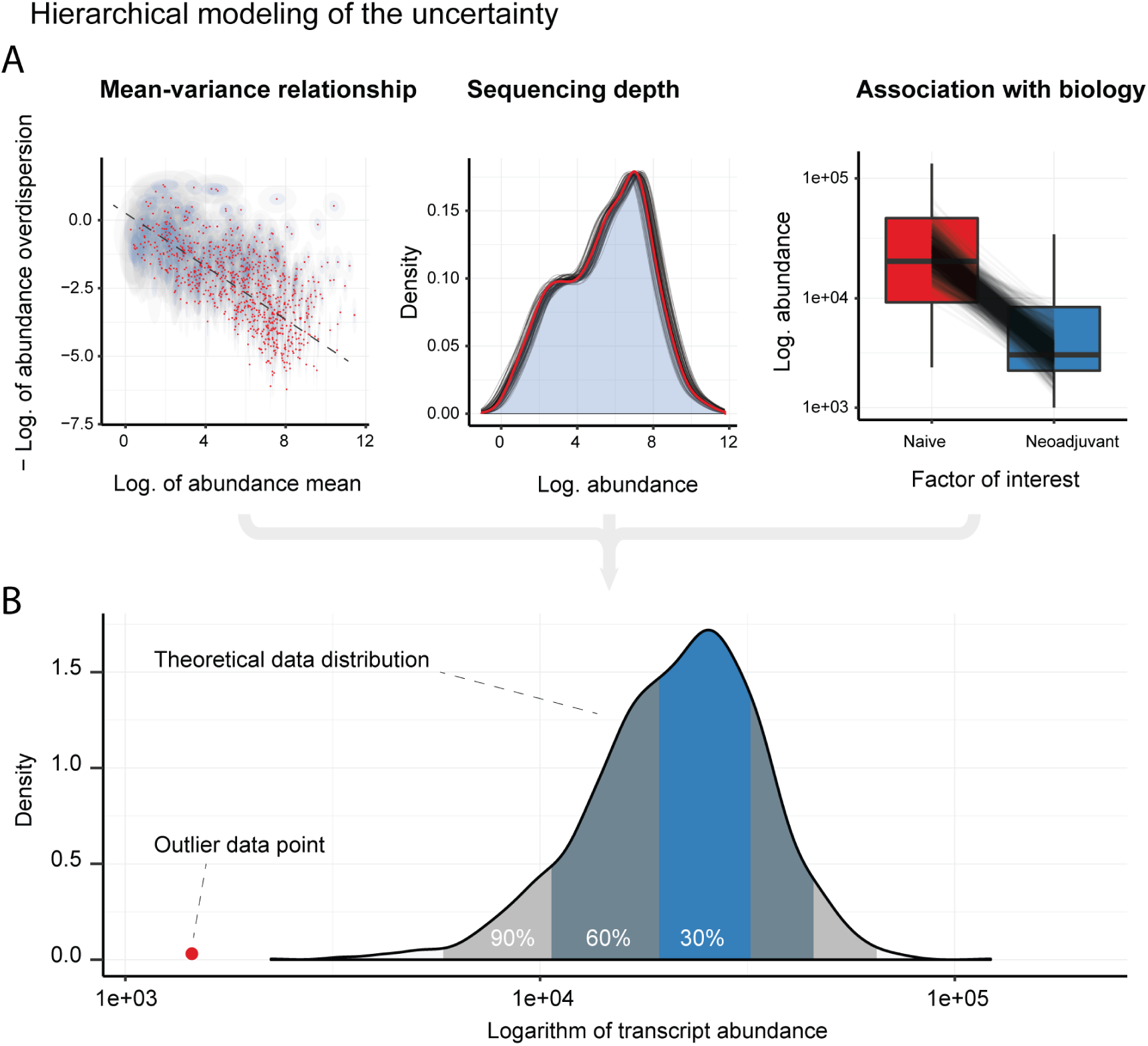
Conceptual map of the modeling of the uncertainty for each transcript/sample pair. **A -** From the left are represented: (i) the uncertainty of mean and overdispersion of each transcript, and their relationship, the red dots represent the point estimates, while the ellipses represent the two-dimensional credible intervals 40% (blue) and 95% (grey); (ii) the uncertainty of the sample-wise effect of sequencing depth, where the overlapping curves represent the possible densities (95% credible interval) of whole transcriptome and the red curve is the mean inference; and (iii) the uncertainty of the association between transcript abundance and the covariates in the generalised linear model, where the overlapping slopes represent the possible associations (95% credible interval) between transcript abundance and factor of interest. **B** - The theoretical data distribution for a transcript/sample pair, resulting from the integrated uncertainty.

**Figure 2.**
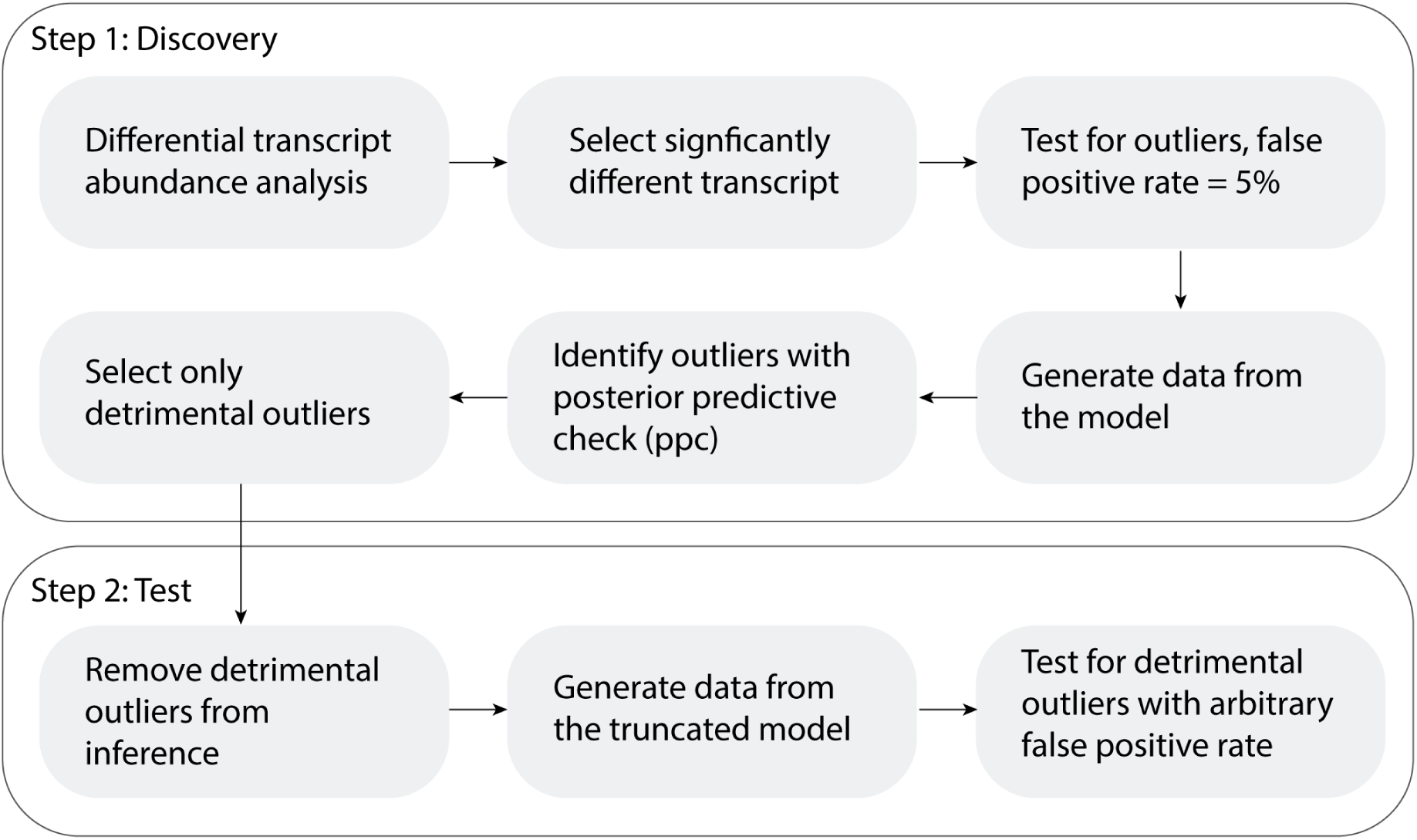
Flow chart of the two-step approach. In the first step, transcript with potential outliers based on the user linear model are identified with the 5% false positive rate. In the second step, a 95% credible interval truncated negative binomial distribution is used, and the transcripts including outliers are identified with a user selected false positive rate.

In the first *discovery* step, the model is fitted to a list of previously identified differentially abundant transcripts. New data is generated from the fitted model, providing the theoretical range of values for each sample-transcript pair. All observed read counts that are outside the 95% posterior credible interval are quarantined as possible outliers. In the second *test* step, the model is fitted again excluding the deleterious outlier data points that would skew the estimated difference between conditions to be larger (i.e. only the combinations (i) higher than the upper quantile of the credible interval when the condition is estimated to have increased expression; or (ii) smaller than the lower quantile when the condition is estimated to have decreased expression) using a truncated negative binomial distribution at 2.5% and 97.5% quantiles (Fig. S1). New data are generated from the second fitted model, and all the observed read counts (including possible deleterious outliers quarantined from the inference) are tested against these, using a credible interval that matches the user-selected false positive rate, assuming the remaining data is generated by a pure negative binomial process. Given the desired false positive rate (1% by default), the interval width is taken as 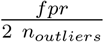 where the factor of two, compensates for unidirectionality of the tests (just for deleterious outliers).

### Probabilistic framework

A Bayesian inference probabilistic network is used to model the raw read counts, based on a negative binomial regression. The differences in sequencing depth across samples are modelled with a sample-wise exposure rate term, that multiplies the transcripts expected abundance. The expected abundance across genes is modelled as generated by a gamma distribution. The over-dispersion is modelled in a gene-wise manner. The inverse association between log transcript abundance and its negative log over-dispersion is modelled as a linear function with a negative slope.

This probabilistic model can be represented by a joint probability density formula (Eq. 1) and a series of sampling statements (Eq. 2–7; Fig. 3). The term *X* represents the design matrix, *α* represents the covariate factors, *ε* represents the exposure rate (sequencing depth effect), *σ* represents the over-dispersion, *μ* represents the mean of the gamma prior for expected transcript abundances, represents variance of such prior, , and Φ represent the intercept, slope and standard deviation of the negative log variance-log mean association.

**Figure 3.**
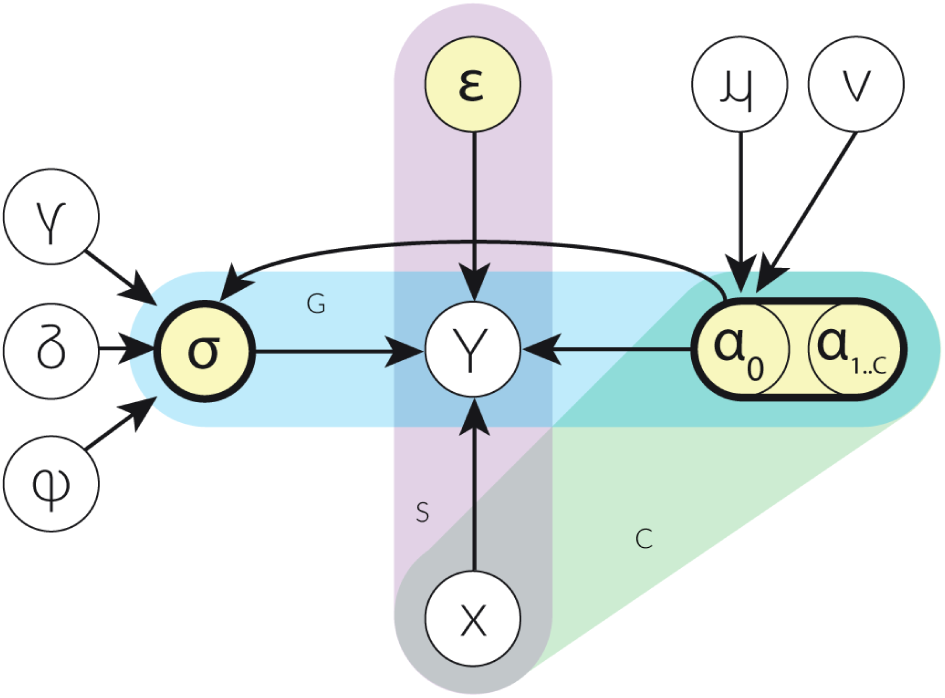
Graphical plated representation of the inference model representing Eq. 1–7. The white circles represent observed data. The yellow circles represent parameters (reals, vectors or matrices). The coloured frames group variables into a subgraph that repeats for transcripts/genes (G), samples (S) and covariates (C).

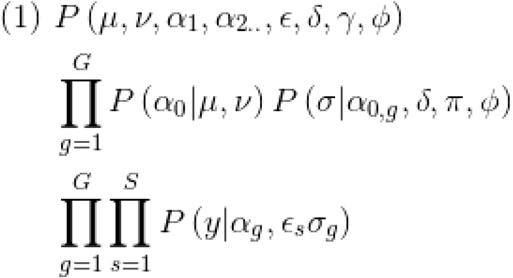

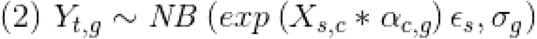

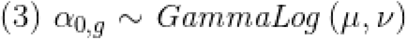

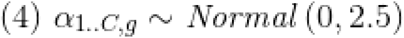

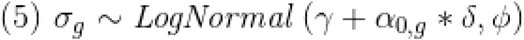

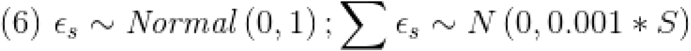

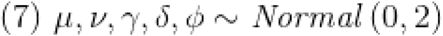

The exposure rates are inferred using a set of anchor genes selected from the bottom rank of differentially abundant gene transcripts provided by the user (n = 500 by default). The underlying assumption is that those genes are not associated with the factor of interest. Internally, the model is parameterised in logarithmic space for higher numerical stability. For the direct calculation of the credible intervals from the generated posterior distribution for the second inference step, the number of generated data drawn *D* from the model is chosen as sufficient to produce on average *N* draws (*N* = 10 by default) outside the confidence interval of choice. The number of total draws needed is calculated as follows.

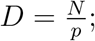

where *p* = 0.05, for the inference discovery step, 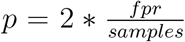, for the inference testing step, N is the desired number of draws outside the credible interval, and *fpf* is the desired false positive rate. For example, if we desire 50 draws outside the credible interval, a false positive rate of 0.1, and the dataset includes 10 samples, we need to draw 2500 points from the posterior distribution. From a theoretical perspective, in case the data was strictly generated from a specific probabilistic model, these example values will provide 1% false outlier classifications. Given 100 genes to be tested, we can expect to obtain one false positive; while if we tested one gene, and repeated the analysis 100 times, we can expect to obtain one false positive overall. The multiplication by two for the testing step is to reflect that we are doing a one tail observation only (testing for deleterious outliers only).

For the second *test* step, the negative binomial distribution of transcript abundance is modelled as truncated at the quantiles 0.025 and 0.975 (corresponding to a 95% credible interval used to quarantine outlier data-points in the first step). Given the computationally demanding calculation of probability densities of a truncated negative binomial, we employ a heuristic adjustment to the over-dispersion of the generated quantities for the calculation of the predictive posterior credible intervals. That is, given that the under-estimation of over-dispersion that a standard negative binomial has compared to a truncated negative binomial at the 95% percentile is approximately constant for any combination of mean and over-dispersion ranges typical in RNA sequencing data (Fig. S1), we use the posterior distribution of mean and over-dispersion from a non-truncated negative binomial and then generate data adjusting the over-dispersion after fitting.

### Posterior probability distribution sampling and approximation strategies

In order to infer and sample from the joint posterior distribution of all parameters, the Bayesian probabilistic framework Stan was used (Carpenter *et al.*, 2017). To optimise its execution, it is possible to approximate the posterior distribution generation using variational Bayes, as opposed to generating the posterior probability distribution using the Hamiltonian Markov-chain Monte Carlo sampling (Neal and Others, 2011). The number of draws needed to explore the distribution tails to identify the credible interval of the predictive posterior distribution increases *quasi*-exponentially with the decrease of false positive rate or the increase in the number of samples. To reduce the computational burden, approximation of the calculation of the credible interval of the theoretical data distribution may be necessary. For these cases, the credible interval of each transcript/sample pair can be calculated analytically using the posterior distribution of the mean, over-dispersion and exposure rate parameters from each sample-transcript pair. Briefly, we sample with replacement the posterior distribution N times for those three parameters, where N is the number of desired samples to define the credible interval accordingly with the user-selected false positive rate; then we generate N random numbers from a negative binomial distribution (R Core Team, 2013).

### Calibration and accuracy test

In order to test the accuracy of the outlier inference, we produced simulated data from the joint posterior distribution fitted on real data (Mangiola *et al.*, 2019), including 339 transcripts to be tested (result of edgeR analysis; FPR < 0.05) across 21 samples. Briefly, we performed edgeR analysis of this dataset and identified potential differentially abundance transcripts (false discovery rate < 0.05) according to a linear model including risk as the only covariate. Those transcripts were modelled with our Bayesian inference model, and the posterior distribution was used to generate simulated data that comes from a pure negative binomial generative process and has all the biological and experimental properties of the source experimental data set. For a random selection of 20% of those transcripts, we injected one outlier for one randomly selected sample, characterised by a 10^−10^ quantile distance from the true posterior of the selected transcript/sample pairs.

We then used these simulated data sets to calculate the false positive and false negative rate testing for 18 user-selected false positive rate thresholds, ranging from 0.2% to 10%, replicating each run three times for a total of 54 runs. We then calculated (i) the proportion of transcripts labeled as containing outliers and compared them with the nominal false positive rate threshold; and (ii) the false negative calls per each nominal false positive rate threshold.

## Results and discussion

### Model calibration

Our test on simulated data showed that the model is well-calibrated for false positive rate (Fig. 4A). The correlation across runs with a wide range of false positive rate thresholds (from 0.001 to 0.1) is close to 1 with a R-square of 0.95. The false negative rate for outliers outside credible interval is 0.37 for an aimed false positive rate of 5%, tested against 339 genes across 21 samples (for a total of 7119 inferences; Fig. 4B and 3C).

**Figure 4.**
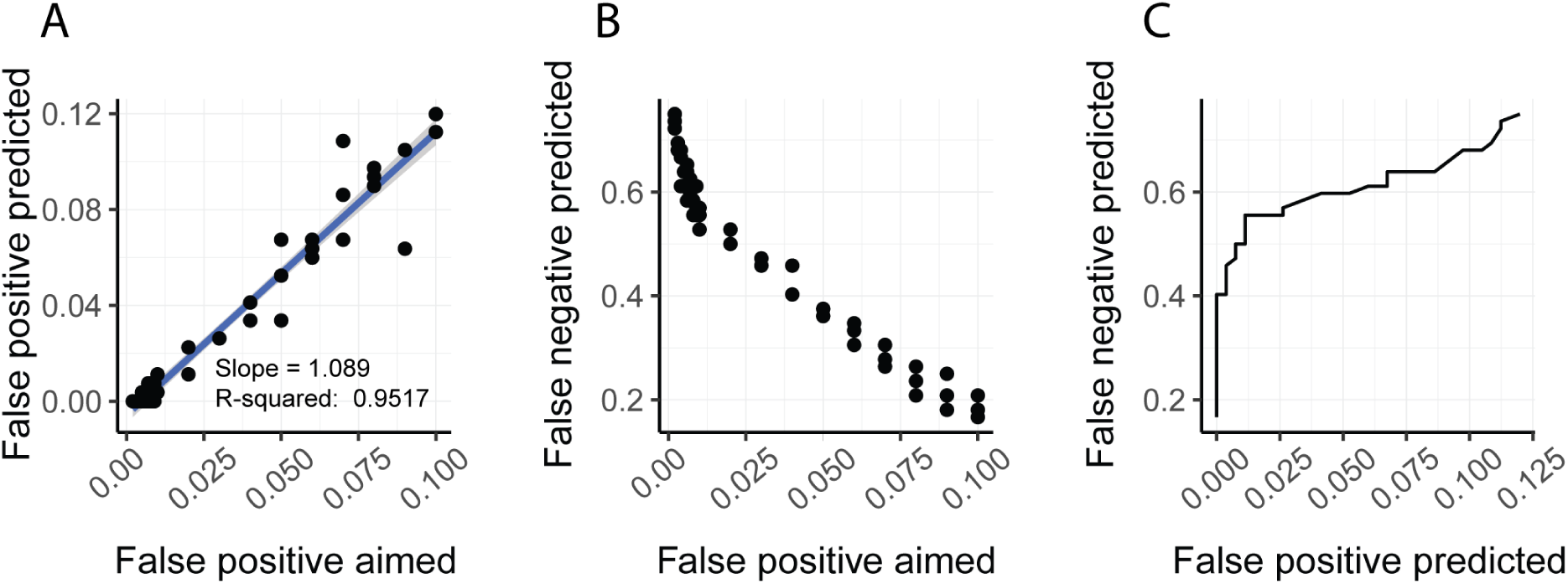
**A** - Scatter plot showing good calibration of false positive rate; representing the linear association between the user defined false positive rate and the false positive rate that the model identified on a simulated data set with no outliers. The statistics are relative to a linear interpolation of the data using the lm function in R. **B** - Scatter plot showing the decrease of false negative with the increase of false positive aimed. **C** - The receiver operating characteristic curve (ROC) of the test analysis. The data points include only inference with the false positive aimed within a meaningful range for standard applications (from 0.002 0.1).

Although our model is well-calibrated against data generated from a negative binomial process, care is needed into making claims about probabilities. In the first discovery step we quarantine data based on the 95th percentile, although this interval is an estimate, given that the presence of outliers makes the numerical generative process not strictly negative binomial by definition. In the second test step, the modeling of the data without quarantined points allows a much better approximation of the a posteriori probabilities and the false positive rate. For the approximation of a truncated negative binomial, we observed that a non-truncated negative binomial under-estimates the over-dispersion for data truncated at the 95ft percentile to an approximately constant degree. The over-dispersion parameter (with over-dispersion being *e*) has a 74% reduction across all mean/sigma combinations that are typical of RNA sequencing data (Fig S1).

### Application to real data, user interface and generated graphics

The application of our model to a dataset of periprostatic tissue (Mangiola *et al.*, 2019) including 21 samples revealed that 9.5% of the differentially abundant transcripts estimated by edgeR had inflated statistics caused by the presence of outliers.. For example, the second top differentially abundant transcript CYP1A1 characterised by an apparent increase in abundance for the neoadjuvant treated group, includes an outlier sample. Its presence noticeably affects the estimation of differential abundance, providing an apparent change of 194 folds, compared to 2.4 if the outlier is excluded. This aspect is visually represented (Fig. 5) by the difference between the dashed credible intervals (inferred including the potential outlier; with 5% false positive rate) and the solid credible intervals (inferred on a truncated distribution, which omits potential outliers; with 1% false positive rate). Figure 4B shows that although the hypothesis test for the transcript CYP1A1 must be run without sample 11165PP, the differential abundance could potentially reach significance, with a much lower fold change. An interesting case is transcript GBP5, for which sample 11184PP is inferred to be an outlier with a raw transcript abundance outside the upper quantile of its credible interval, although it is not characterised by the highest abundance across all samples overall. This inference is mostly driven by the low sequencing depth (proportional to the size of the dots) of the sample 11184PP, which does not justify the relative high abundance for this transcript.

**Figure 5.**
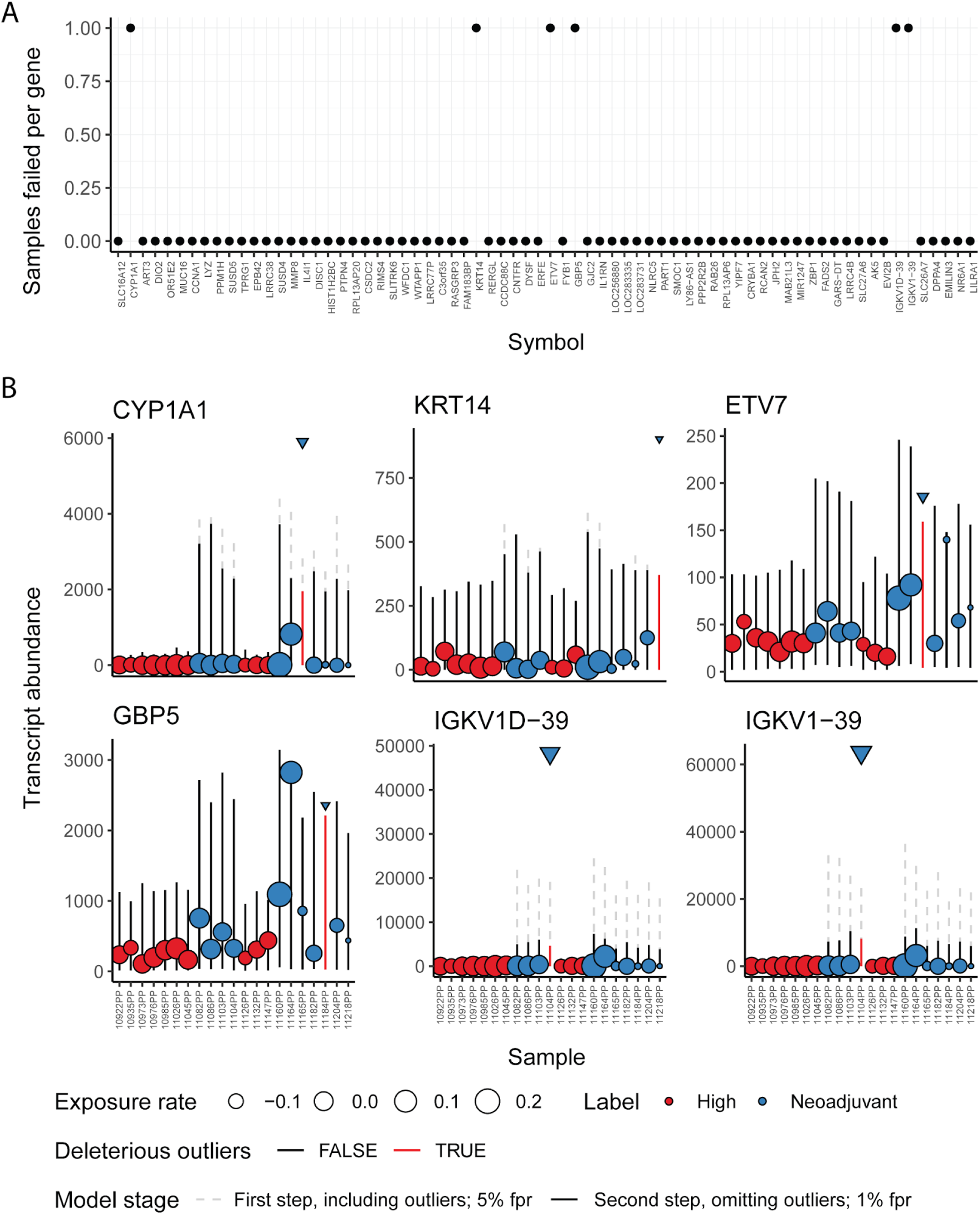
**A** - Dot-plot representing the number of outliers detected in the top differentially abundant transcripts inferred by edgeR on a public data set (Robinson *et al.*, 2010; Mangiola *et al.*, 2019). B - Visualisation produced by ppcseq R package of the 6 transcripts from panel (A) including outliers that inflated the edgeR statistics. The color coding represents the treatment regime, the error bars represent the credible interval of the theoretical data distribution, the size of the points is proportional to the inferred sequencing depth factor (exposure rate). The dashed error bars represent the 95% credible interval of the theoretical data distributions including outliers (first discovery stage), while the solid error bars represent the 99% credible interval (user defined parameter) data distribution excluding outliers, derived from a truncated (at 95ft percentile) negative binomial distributions. The red error bars and the triangular data points represent the outlier observations that do not fit the model.

The centrality of the use of an iterative approach including a truncated, distribution is supported by attempts to identify outlier data points with only one passage (i.e. discovery stage) with an approximate false positive rate of 1%. Using this false positive rate no outliers could be detected, mainly because the presence of deleterious outliers significantly inflates the change in transcript abundance between the two conditions.

### Efficient approximation of posterior probability density and credible interval provide comparable accuracy to Hamiltonian Monte Carlo

The test runs performed with increasing level of parallelisation (from 2 to 16 physical cores) show a gradual speedup to 3 times that of the non-approximated model (Hamiltonian Monte Carlo sampling with calculation of quantiles through posterior draw; Fig. 2S). Compared to the non-approximated approach, the variational Bayes approximation showed speedup from 2- to 6-folds depending on the level of parallelisation of the alternative Hamiltonial Monte Carlo sampler (from 16 to 2 physical cores respectively; Fig S2).

As the amount of draws from the posterior probability distribution needed to define extreme quantiles of a distribution grow exponentially with the number of samples and the false positive rate, a further approximation of the calculation of the quantiles is needed for practical purposes (both for Hamiltonian Monte Carlo and Variational Bayes). The accuracy compared to the ground truth is high (Fig. 6A), with a relative error of the distribution mean (average across all approximation combinations) of 0.10, a relative error of the lower quantile of 0.04 and of the upper quantile of 0.71. The approximation combinations do not affect the inference compared with the non-approximated approach (Fig. 6B), and bias in the under-estimation of the negative binomial variance is not noticeable. Overall, the correction lets us restore almost exactly the full Bayesian posterior intervals while fitting the model with the more efficient variational Bayes approach.

**Figure 6.**
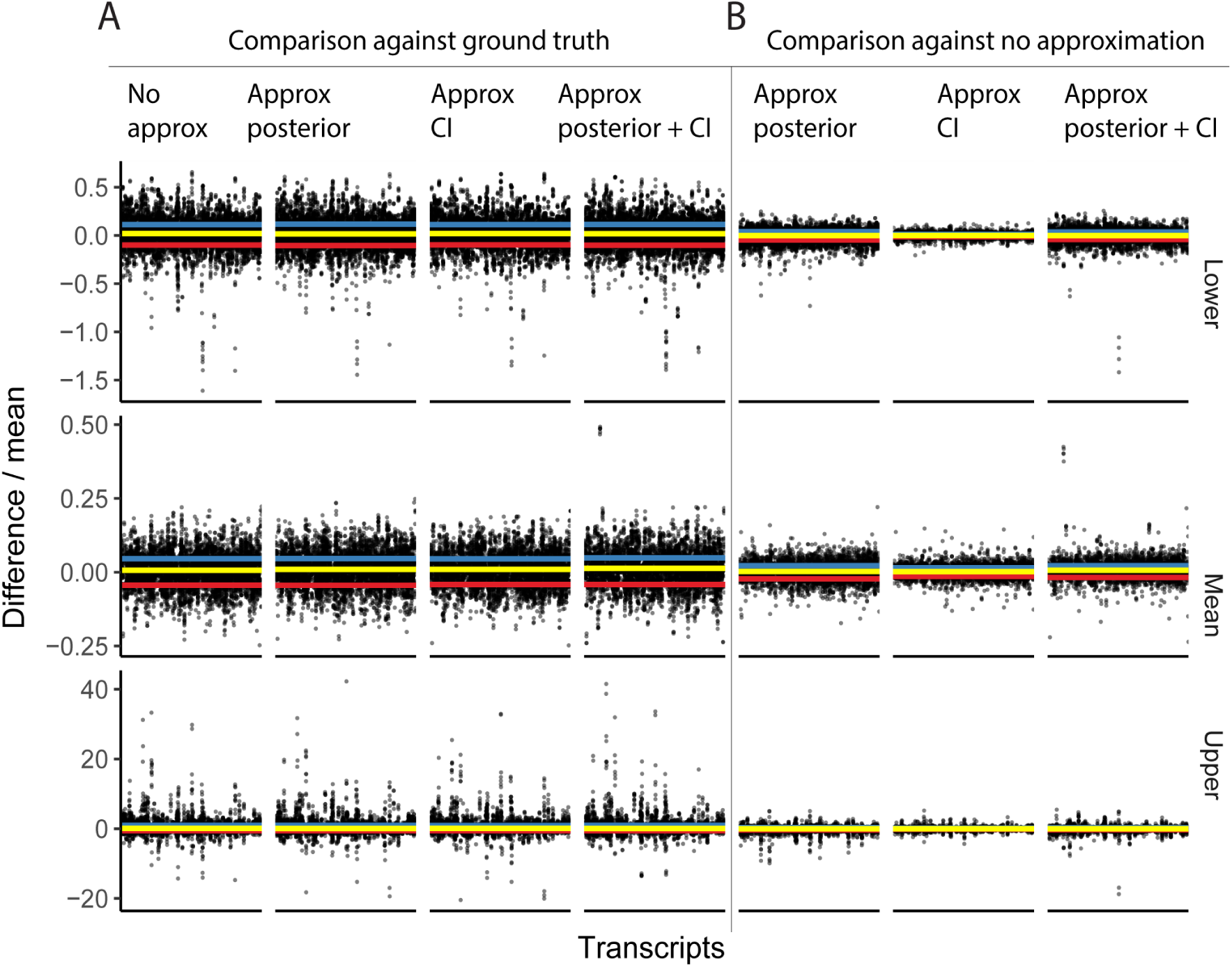
**A -** Relative error of the mean, lower and upper quantiles across, compared to simulated data based on a regression model from the real data set (Mangiola *et al.*, 2019). **B -** Relative error of the mean, lower and upper quantiles across, compared to the non approximated modeling. The credible intervals 95%, 99%, 99.5% and 99.9% were tested. The yellow horizontal line corresponds to the median error, the blue and red lines correspond to the upper and lower standard deviation.

## Conclusions

Differential transcript abundance analyses are key in many areas of biology, and often studies include a limited number of biological replicates. In these cases, the effect of outlier observations can have a disproportionate impact on the prioritisation of differentially abundant transcripts. Therefore, it is important to be able to quarantine transcripts for which the statistics are driven by observations that do not fit the model assumptions. It is possible to identify outlier observations by analysing the distribution of residuals; however, in cases where limited biological replicates are available this analysis tends to be under-powered. The use of Bayesian inference allows a posterior predictive check, where the theoretical range of values for each observation is estimated by sharing the uncertainty across transcripts (e.g. the association of mean and over-dispersion) and samples (the sequencing depth unwanted variation).

Here, we propose a robust statistical framework for the detection of transcripts for which data does not fit the assumption of a negative binomial distribution, including deleterious outliers that bias the statistical inference toward false positives. This process includes two steps, where transcripts for which the statistics are biased by potential outliers are flagged and the likelihood of this event is calculated based on a truncated distribution, which helps to control false positives. The user is able to control for an arbitrary rate of false positives at the transcript level, which is a direct and intuitive measure of confidence. This method can be used to check and visualise results from all differential transcript abundance methods based on a negative binomial framework (e.g., edgeR and Deseq2) providing a more robust differentially abundant transcript set.

## Online methods and raw data

The code used to conduct the analyses is available at github.com/stemangiola/ppcseq.

## Acknowledgements

We thank Dr. Damiano Spina (RMIT) for the support with machine learning.

## Authors contribution

SM conceived and designed the method under the supervision of ATP. MM contributed to statistical model implementation and checking. EAT contributed to benchmarking analyses. All authors contributed in manuscript writing.

## Supplementary Figures

**Fig S1.**
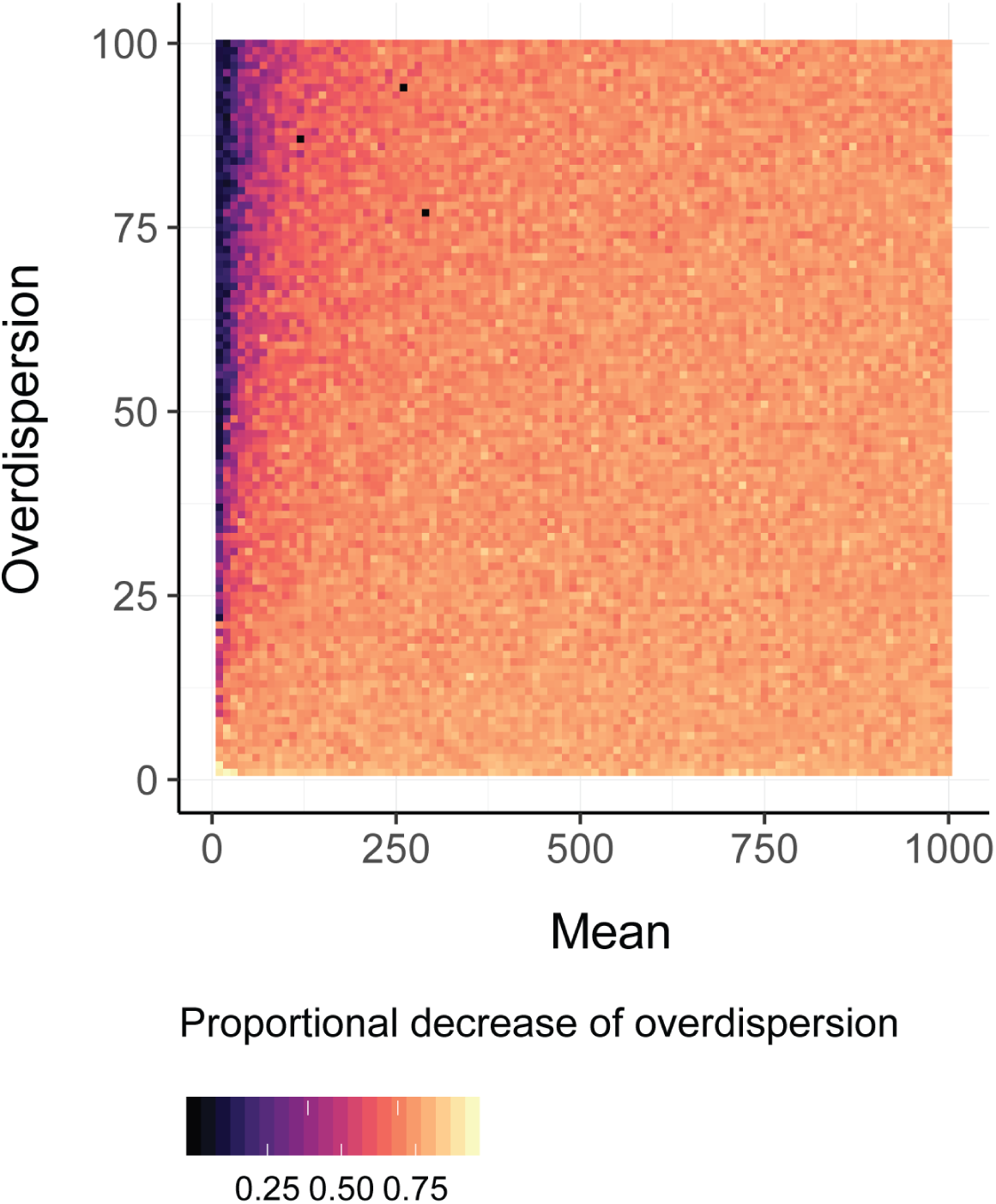
Proportional decrease of over-dispersion of a negative binomial following a truncation of the distribution at the 2.5% and 97.5% percentiles.

**Fig S2.**
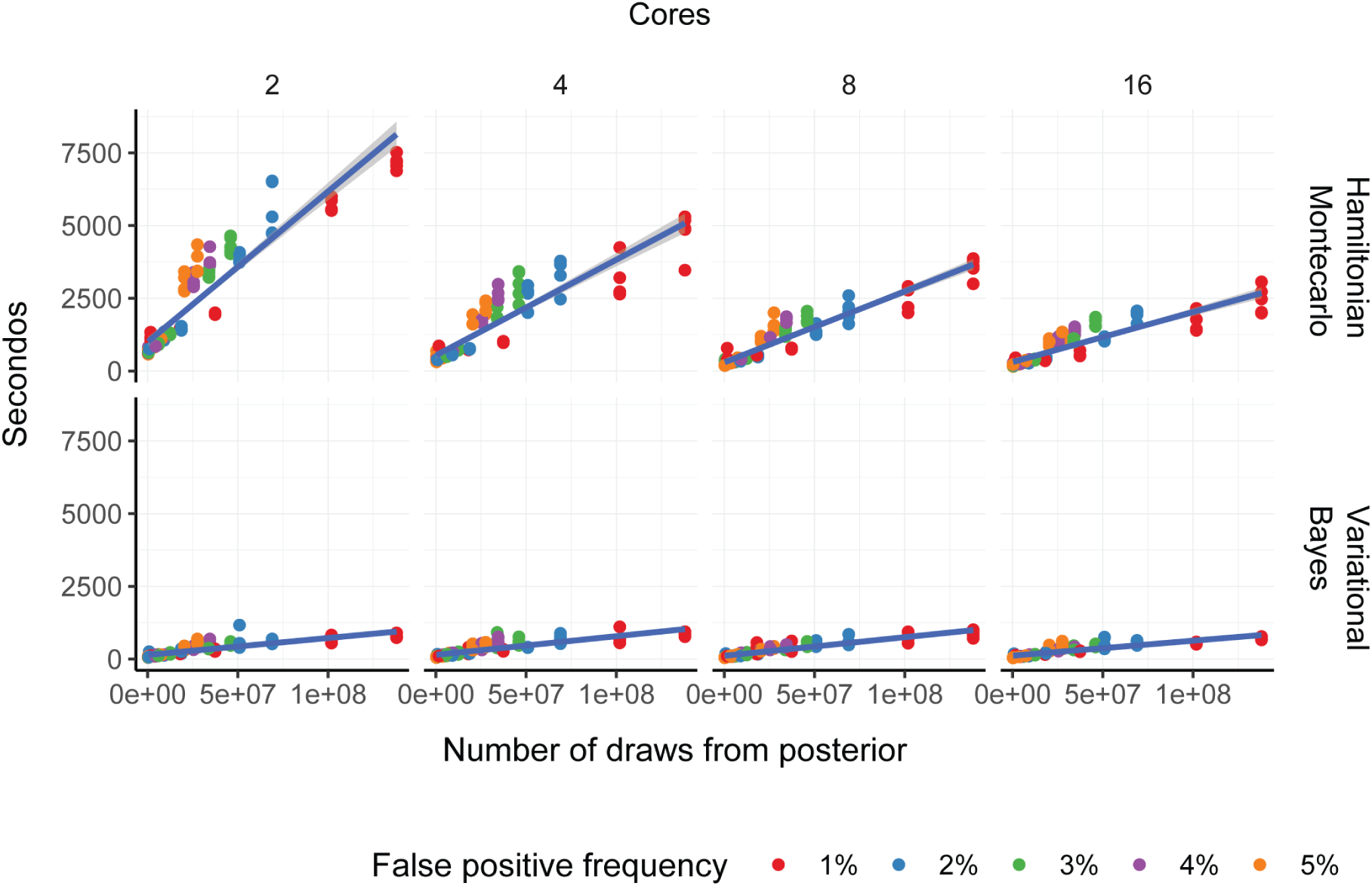
Benchmark of execution time across runs with diverse user-defined false positive rates, between full posterior sampling using Hamiltonian Monte Carlo and variational Bayes approximation (multivariate normal).

**Fig. S3.**
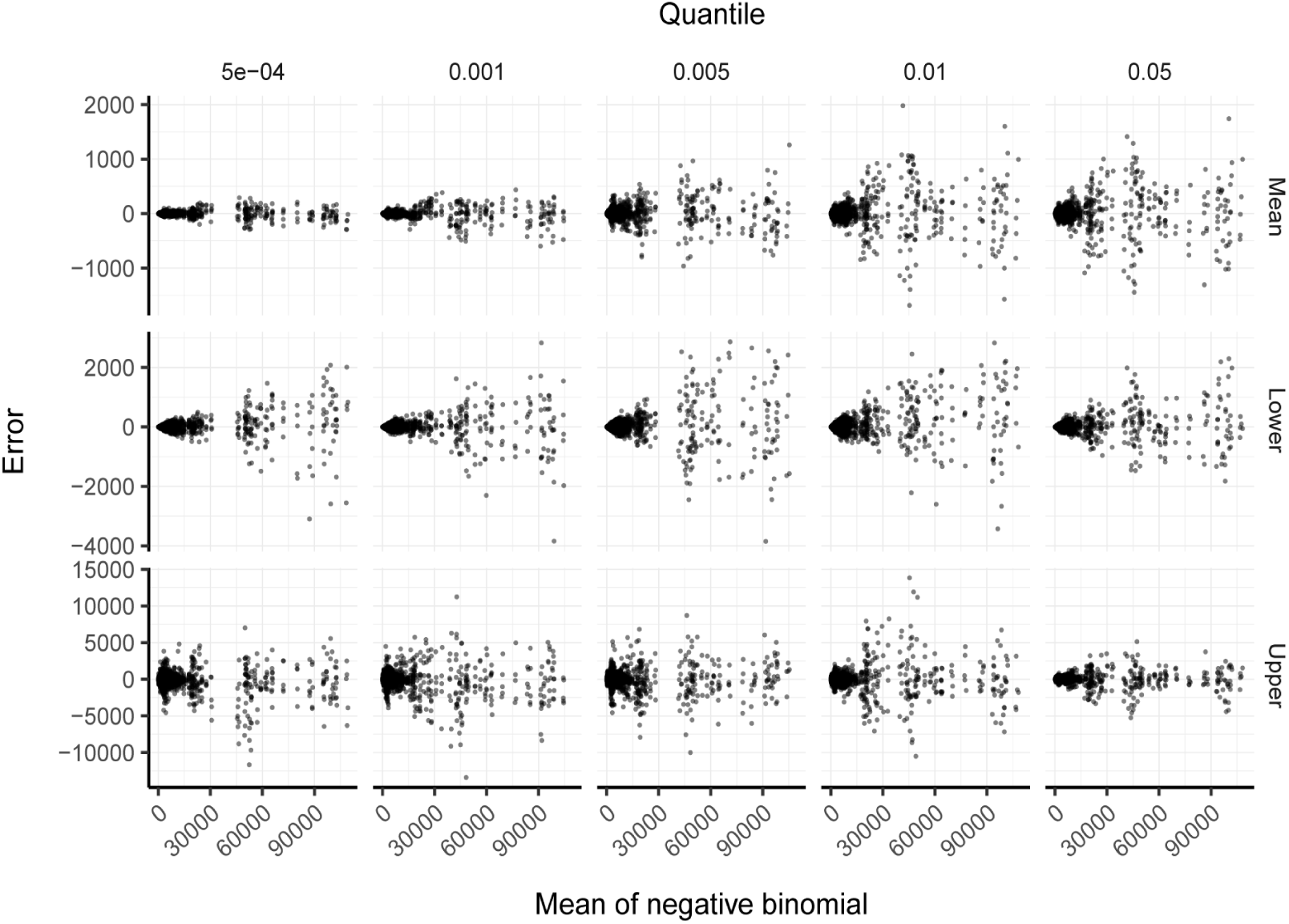
Association between difference between credible interval estimation using posterior draws (standard procedure) and using analytical approximation against mean before and after adjustment (see Materials and Methods). This figures shows the absence of bias of credible interval estimation using the approximation method compared with the pure calculation based on posterior probability draws.

